# An Antagonistically Pleiotropic Gene Regulates Vertebrate Growth, Maturity and Aging

**DOI:** 10.1101/2023.05.01.538839

**Authors:** Eitan Moses, Roman Franěk, Tehila Atlan, Henrik von Chrzanowski, Elizabeth Duxbury, Shay Kinreich, Alexei A. Maklakov, Itamar Harel

## Abstract

The antagonistic pleiotropy theory of aging predicts functional trade-offs between early-life and late-life fitness. However, empirical evidence for these trade-offs in vertebrates remains scarce, particularly in the context of ecologically relevant life histories. Here, we identify *vestigial-like 3* (*vgll3*), a transcription cofactor previously linked with age at maturity in humans and Atlantic salmon through GWAS studies, as an antagonistically pleiotropic gene in turquoise killifish (*Nothobranchius furzeri*). By disrupting two conserved *vgll3* isoforms, we show that reduction of *vgll3*, in an isoform- or dose-dependent manner, accelerated male growth and reproductive development. This indicates that *vgll3* regulates sexual maturity. However, early-life benefits come at a late-life cost, as older mutant males with a disrupted long isoform develop melanoma-like tumors, validated via transplantation into immunodeficient *rag2* models, and exhibit increased age-related mortality rate. These findings highlight *vgll3* as a key regulator of vertebrate life-history trade-offs, balancing early-life fitness with late-life disease risks.

## Introduction

Vertebrates exhibit a remarkable degree of variation in their life history traits, including size, age at maturity, and lifespan. This extreme diversity is exemplified by the ∼1000-fold difference in lifespan between the short-lived turquoise killifish^1^ and the centuries-old Greenland shark^2^. The antagonistic pleiotropy theory of aging^3^ (AP) aims to explain this variation, by proposing that some genes may enhance fitness early in life while being detrimental late in life, leading to a positive selection of genes that ultimately limit lifespan^4–10^.

Insight into the regulatory networks that may include AP alleles can be gained, for example, through genome-wide association studies (GWAS) that link early-life performance and age-related diseases^11–17^. Other approaches, such as experimental evolution in model organisms, have demonstrated that selection for late-life reproduction can increase lifespan and reduce early-life fecundity^18–20^. However, such studies do not address the underlying genetic mechanisms at a single gene resolution, which could be determined through genetic perturbations (e.g. the *C. elegans* trl-1 mutant^21^).

Several targeted gene mutations with increased longevity have been identified in laboratory animals^22^, including mice. Some of these mutations were shown to have robust trade-offs with growth and reproduction, as seen in the Ames and Snell dwarf mice^23–26^, and following genetic manipulation of IGF1 signalling^27,28^. However, there are few examples in vertebrates where putative molecular mechanisms of antagonistic pleiotropy are directly linked to the regulation of specific life-history traits within a normal, biologically relevant range of expression (but see^24^). For instance, Ames and Snell dwarf mice are long-lived but sterile, indicating that such deleterious mutations are unlikely to contribute to natural variation in lifespan and aging within wild populations.

Recent GWAS in wild salmon populations^29,30^ and humans^31,32^ link polymorphism in the vestigial-like family member 3 (*vgll3)* locus, with a variation in age at maturity in the wild. These findings suggest VGLL3 might have evolutionarily conserved functions regulating pubertal timing in vertebrates. Fish species exhibit a vast range of evolutionary adaptations, providing an experimental playground for modeling complex traits in health and disease^22,33–36^. Here, we leverage the turquoise killifish (*Nothobranchius furzeri*) as a model to investigate the role of VGLL3 as a regulator of antagonistic pleiotropy in vertebrates.

With their rapid sexual maturation (∼2–3 weeks in the wild^37^) and a naturally short lifespan (4– 6 months^1^), killifish present an excellent platform for experimental manipulation of vertebrate life history. By targeting two evolutionarily conserved *vgll3* isoforms, we reveal that their disruption accelerates male somatic growth and advances puberty onset in a dose- and isoform-dependent manner. Remarkably, aged mutants exhibit melanoma-like tumors with high penetrance, validated through cancer engraftment studies into a newly developed immunodeficient *rag2* mutant killifish. Lifespan analyses indicate that male *vgll3* mutants also exhibit a greater increase in mortality rate with age. These findings establish *vgll3* as an antagonistically pleiotropic gene, mediating a trade-off between early-life benefits, such as accelerated growth and reproductive maturity, and the heightened risk of late-life disease.

## Results

### Cell-type and sex-specific gonadal expression of *vgll3*

In humans, VGLL3 protein expression is primarily found in the male and female gonads^38^. By analyzing single-cell data that we have recently produced from the killifish testis and ovary^39^, we identified which specific gonadal cell types express *vgll3* (**Figures 1a**, left, **and S1a**). Our analysis indicated that *vgll3* is primarily expressed in the somatic cells of the gonad, particularly in male-specific Sertoli and Leydig cells (**Figure 1a**, right, and **Figure S1b**), highlighting its possible sexually dimorphic role.

**Figure 1:**
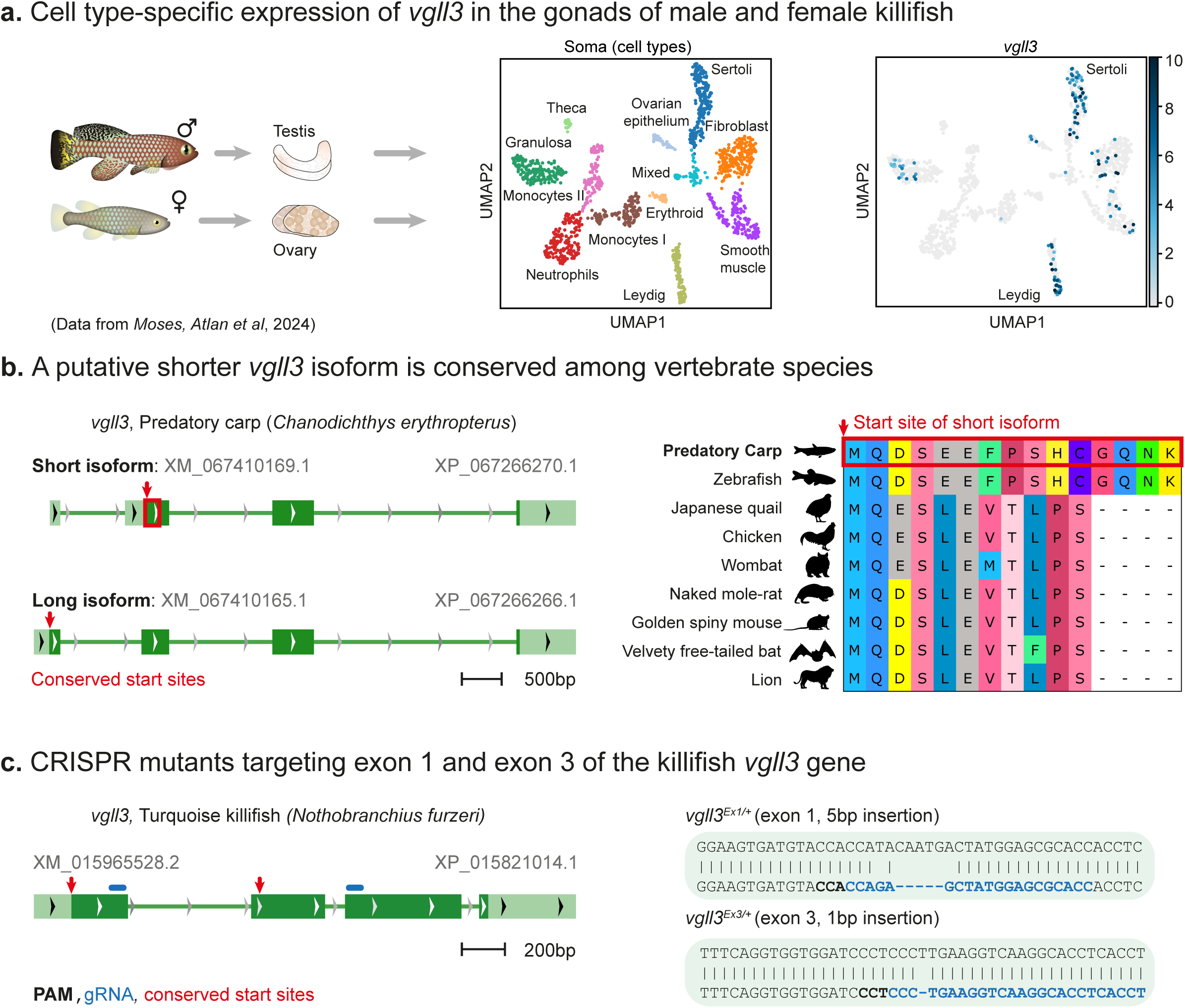
Generation of two *vgll3* mutants, corresponding to conserved isoforms. **a)** UMAP analysis of single cells from the testis and ovary of 1-month-old fish (adapted from^39^). Somatic cells are clustered, and color-coded by cell types (center). UMAP plot of *vgll3* expression in specific cell types (right). **b)** Gene model of both long and short isoforms of the *vgll3* gene in the predatory carp (*Chanodichthys erythropterus*). Red arrows indicate the location of the start site of the short isoform in both gene models (left). Multiple alignments of the start site of the short isoform in various vertebrate species, red box marks the corresponding reading frame marked in the left panel (right). **c)** Generation of two *vgll3* CRISPR mutants corresponding to both isoforms. A gene model of the killifish *vgll3* with red arrows indicating the start sites of both isoforms, and blue lines indicating the gRNA targets (left). Specific DNA sequences from the long and short isoforms with gRNA sequence (blue), protospacer-adjacent motif (PAM, in bold), and indel (right).

We then investigated the spatial distribution of the *vgll3* transcript in male gonads using single-molecule fluorescence in situ hybridization chain reaction (smFISH HCR, **Figure S1c**). To identify specific cell types, we compared *vgll3* expression with known markers, including *amh* (anti-Müllerian hormone) for Sertoli cells, and the classical germ cell marker *ddx4/vasa* (dead-box helicase 4) as a reference. Our findings indicate that *vgll3* is enriched in male gonadal support cells.

### Genetic manipulation of distinct *vgll3* exons in killifish

Recent findings in salmon suggest that the *vgll3* genotype, associated with either early or late onset of puberty^29,30^, may influence specific *vgll3* transcript isoforms^40,41^. These include a putative shorter isoform in salmon that originates from an alternative 5’ UTR within the first intron, and is likely translated from start codons located in exon 2^40^.

Interestingly, we observed that a shorter conserved *vgll3* isoform is present across many vertebrate species (**Figure 1b**), including fish, such as the predatory carp (*Chanodichthys erythropterus*) and zebrafish (*Danio rerio*). Other vertebrates include birds (e.g., the chicken *Gallus gallus* and Japanese quail *Coturnix japonica*), smaller mammals like the naked mole-rat (*Heterocephalus glaber*) and the golden spiny mouse (*Acomys russatus*), as well as bats (the Velvety free-tailed bat *Molossus molossus*). It also appears in larger mammals, such as the lion (*Panthera leo*), and marsupials like the common wombat (*Vombatus ursinus*). Transcripts mostly originate from an alternative 5’ UTR within the first intron (e.g. carp, **Figure 1b**, left), and are translated from a conserved reading frame (**Figure 1b**, right)

Therefore, to investigate whether VGLL3 and its isoforms have a functional role in vertebrate life-history traits, we edited the killifish *vgll3* using established CRISPR protocols^42,43^ (**Figure 1c**). Specifically, we edited either the first or third exons of *vgll3* (*vgll3^Ex1^*and *vgll3^Ex^*^3^), introducing frameshift mutations predicted to result in loss-of-function alleles (**Figure 1c**, and see **Methods**). Importantly, targeting the third exon is expected to disable both the short and long isoforms (**Figure 1c**). We then outcrossed heterozygous fish for several generations to reduce the burden of possible off-target mutations (see **Methods**). As antibodies for fish VGLL3 are not available, qPCR analysis (targeting *vgll3* exon3) revealed significantly reduced *vgll3* expression levels in mutant-derived primary cells, likely due to nonsense-mediated decay (NMD, **Figure S1d**, and previously reported in other killifish mutations^44^).

### *vgll3* mutants display accelerated growth and maturity

Nuptial coloration, a body-color change associated with permanent or seasonal sexual maturation, is common across various vertebrates and invertebrates, and plays a crucial role in reproductive success. In killifish, nuptial coloration is mostly visible in the male caudal fin and was previously shown to be correlated with sperm production and mating^44,45^. Therefore, as a proxy for sexual maturity, the variability in the onset of tail coloration was visually scored at the age of one month (**Figure 2a**).

**Figure 2:**
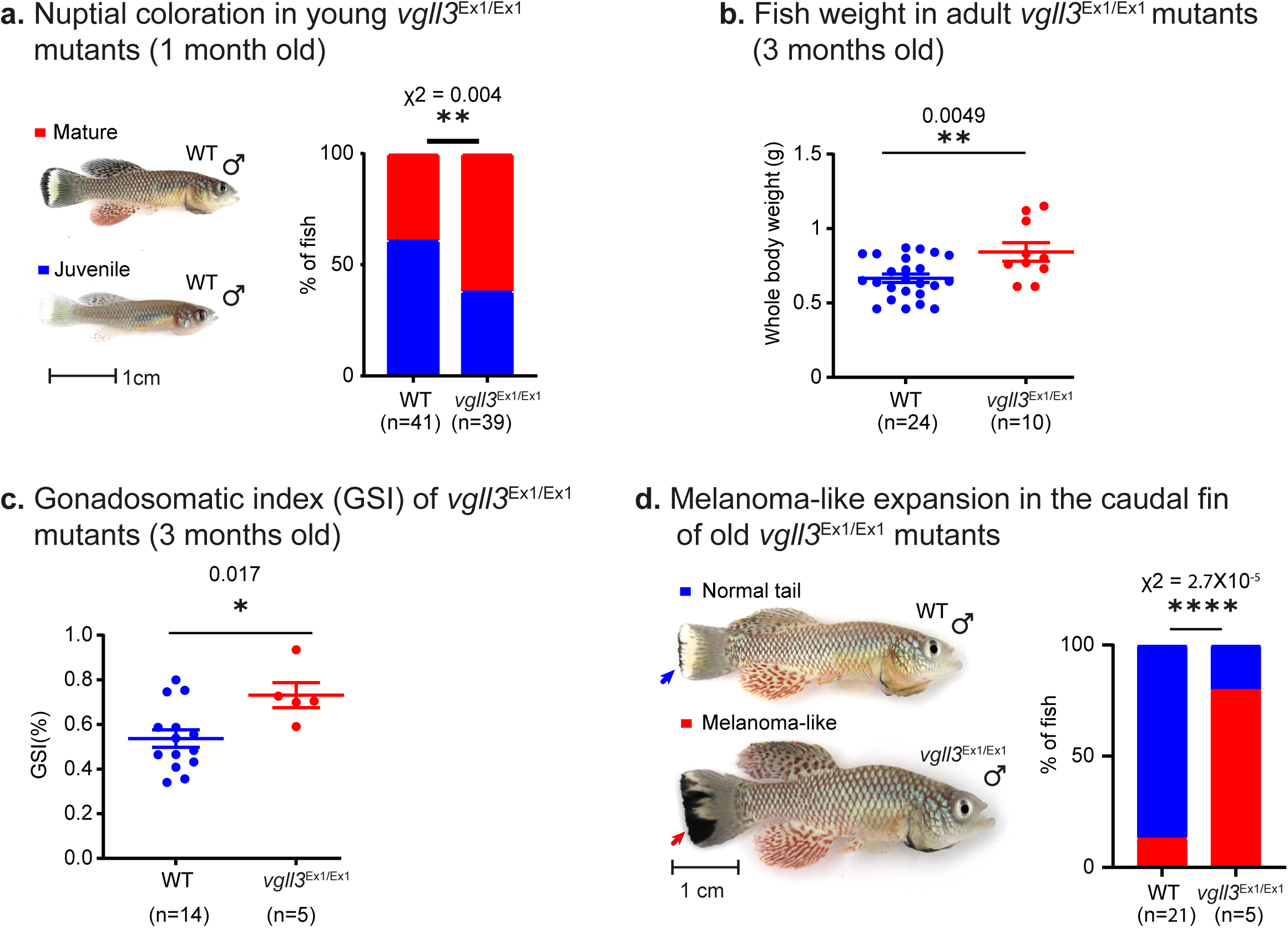
Pleiotropic effects of *vgll3^Ex^*^1*/Ex*1^ mutants. **a)** Left: Representative images of adult WT (1-month-old) males, displaying mature (top) and juvenile (bottom) nuptial coloration. Scale bar: 1 cm. Proportion of fish displaying mature and juvenile coloration in 1-month-old male WT and *vgll3^Ex^*^1^*^/Ex^*^1^ fish. Significance was measured using a χ^2^-test with the WT proportion as the expected value. **b)** Quantification of fish weight in three-month-old WT or *vgll3^Ex^*^1^*^/Ex^*^1^ fish. Error bars show mean ± SEM. Significance was measured using a two-sided Student’s t-test. **c)** Quantification of GSI of three-month-old WT or *vgll3^Ex^*^1^*^/Ex1^*fish. Error bars show mean ± SEM. Significance was measured using a two-sided Student’s t-test. **d)** Left: representative images of old WT and *vgll3^Ex^*^1^*^/Ex^*^1^ males (see **Methods**). The mutant male is exhibiting a melanocyte expansion in its tail (red arrow). Scale bar: 1 cm. Right: proportion of old fish displaying large melanocyte expansions in WT and *vgll3^Ex^*^1^*^/Ex^*^1^ individuals. Significance was measured using a χ^2^-test with the WT proportion as the expected value.

Our data suggest that a significant acceleration in male maturation occurs in homozygous 5bp insertion in exon 1 (*vgll3^Ex^*^1*/Ex*1^*, p*=0.004, **Figure 2a**), and heterozygotes mutations in exon 3 (*vgll3^Ex^*^3*/+*^*, p*=0.005, **Figure S2a**). Interestingly, homozygous mutants for exon 3 (*vgll3^Ex^*^3*/Ex*3^) displayed a reverse trend, indicating a dose-dependent role of each isoform (**Figure S2a**). To specifically isolate the physiological role of the putative short isoform, we focused on homozygous *vgll3^Ex^*^1*/Ex*1^ mutants. Maturation was also associated with higher body weight (*p*=0.0049, **Figure 2b**), and fish size (depth) at older ages (p=0.0012, **Figure S2b**).

The gonadosomatic index (GSI) measures gonad mass as a percentage of total body mass, and is calculated with the formula: GSI = (gonad weight / total body weight) × 100. This index is used to directly assess the sexual maturity of animals, as it correlates with the development of ovaries and testes. While both gonadal weight and body weight increased in *vgll3^Ex^*^1^*^/Ex^*^1^ mutant fish (**Figures 2b, S2c**), GSI was significantly higher, further supporting accelerated sexual maturation (*p*=0.017, **Figure 2c**). Accordingly, a similar increase was observed in the relative testicular area populated with mature sperm (*p*=0.000048, **Figure S2c**, right). Since the testis comprises ∼1% of total body weight, its impact on fish weight is negligible.

These findings are remarkable, given that killifish hatchlings already exhibit one of the fastest growth rates among vertebrates, reaching sexual maturity in as little as ∼3 weeks^1,46^. Together. our data suggests that *vgll3* may act as a rheostat of vertebrate maturity.

### *vgll3* mutants exhibit an increase in age-related melanoma-like expansion

Is there a ‘cost’ for accelerating maturity and growth through *vgll3*? The antagonistic pleiotropy theory of aging proposes that certain genes enhance an organism’s performance early in life but later have harmful effects. These genes can be favored by natural selection because the force of selection on traits is stronger during early life^7^, and such genes can be selected for even if they contribute to a decline in late-life performance.

To explore this paradigm, we aged WT and *vgll3^Ex^*^1*/Ex*1^ mutant populations. Interestingly, we observed that older mutant males exhibited a significant increase in melanoma-like expansion incidences in the caudal fin, suggesting a potential evolutionary trade-off (**Figure 2d**). However, confirming whether these melanocytic expansions are truly cancerous requires developing a set of tools for functional validation.

### Generation and characterization of a *rag2* immunodeficient model

Tumor transplantation studies into an immunodeficient host are a vital approach for investigating cancer biology in model organisms^47–49^. As the first step, we generated an immunocompromised genetic model to enable cancer engraftment by mutating the *rag2* gene^49–51^ (**Figure 3a**). Specifically, we generated a ∼250 base-pair deletion allele (Δ250) in the single exon of the killifish *rag2* coding sequence (**Figure 3a**, left). We outcrossed this line for several generations and generated a fertile homozygous fish.

**Figure 3:**
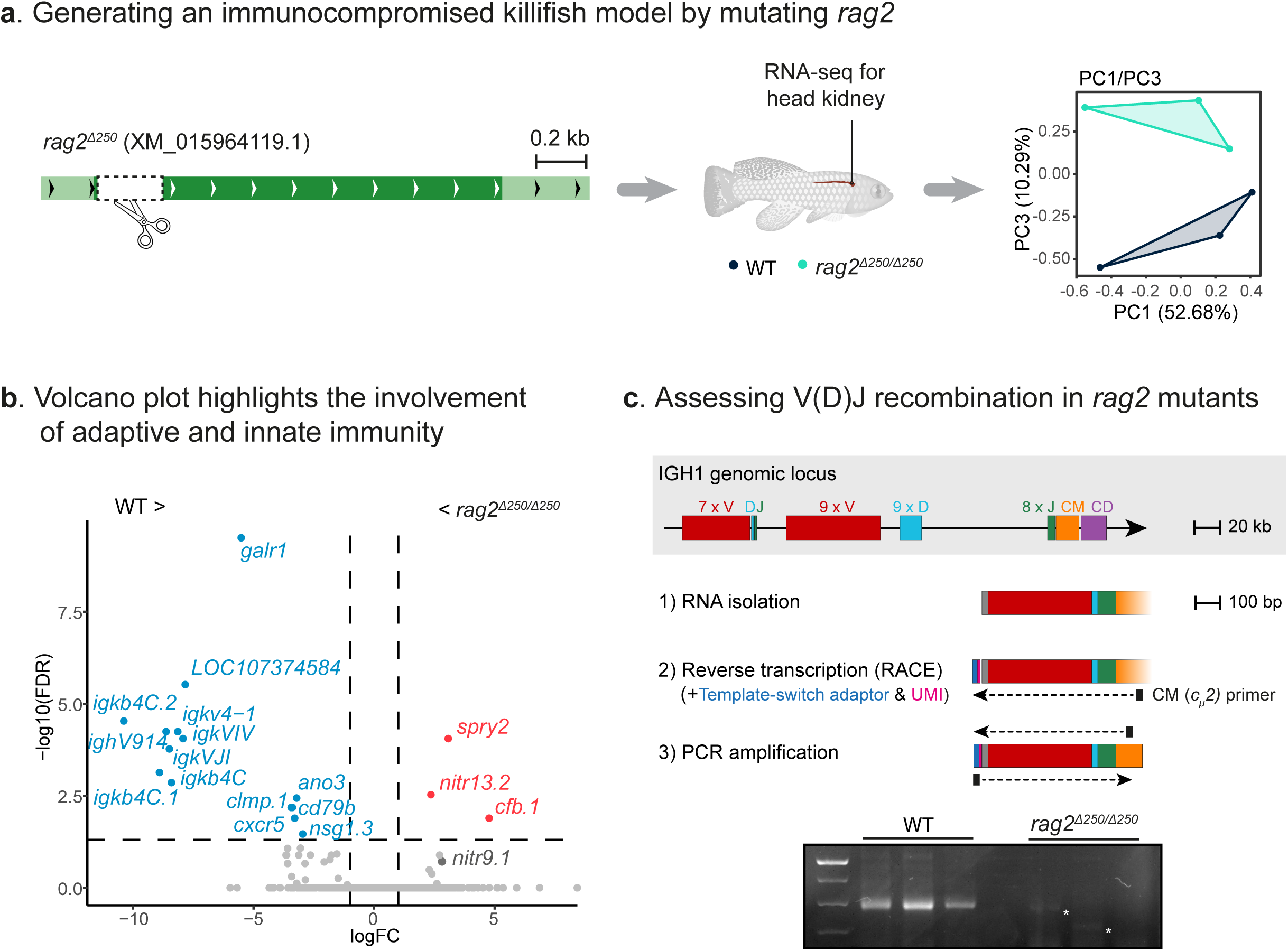
Generation of an immunodeficient killifish model. **a)** Generation of a *rag2* CRISPR mutant with a ∼250 bp deletion (Δ250). A gene model with a single exon is shown (left). Following RNA sequencing for the head kidney of WT (black) and *rag2* mutant fish (green), principal component analysis (PC1/PC3) is shown. Each symbol represents an individual fish (right). n=3 for each genotype. **b)** Volcano plot presenting the differential gene expression between WT and *rag2* mutant fish. The dashed lines represent the statistical thresholds for the GO analysis (fold-change = 2, FDR = 0.05). Upregulated genes are in red, and downregulated genes are in blue. **c)** A schematic model of the killifish *igh* genomic locus, including the constant region (C) and variable regions (top). Experimental design of the V(D)J recombination assay, including RNA isolation, region-specific reverse transcription, and PCR amplification (center, adapted from^62,63^). Agarose gel of the PCR amplified *igh* locus from WT and *rag2* mutant fish, n=3 for each genotype (bottom). Non-specific amplification is marked by an asterisk.

Murine *rag2* mutants display hematopoietic and immune system defects, including arrested B-cell and T-cell development^52^. Therefore, we compared the transcriptome of the killifish head kidney— a dual-function organ responsible for blood filtering and hematopoiesis in teleosts^53^—from three male WT and three male *rag2* mutants. Principal component analysis (PCA) demonstrated that both groups segregate according to PC3 (**Figures 3a**, right, and **S3a**). The differential expression analysis revealed 14 downregulated and 3 upregulated genes (fold change > 2, FDR <0.05, **Figures 3b**, **Table S1**). Interestingly, downregulated genes were directly related to *rag2* functions, including members of the immunoglobulin gene family (**Figure S3b**). These data suggest that failure to rearrange the V(D)J gene segments perturbs expression.

Upregulated genes were sparse, and linked to immune and signaling-related pathways, including the novel immune-type receptors (NITRs)^54–59^, (i.e. *nitr13.2*, **Figure S3b**). NITRs have been identified in all major lineages of teleost fish^60^, and are structurally similar to the killer immunoglobulin-like receptors (KIR) on mammalian natural killer (NK) cells^58^. These receptors do not depend on V(D)J recombination^60^ and are expressed in teleost NK-like cells^58,61^. Interestingly, a similar expansion of NK-like cells was detected in zebrafish *rag2* mutants^57^, suggesting an evolutionary conserved mechanism.

### Lack of V(D)J recombination in *rag2* mutants

The killifish immunoglobulin heavy chain (*igh*) gene locus^62^ consists of a constant region (C), and several variable V(D)J regions (**Figure 3c**, top). To assess V(D)J recombination in *rag2* mutants we adapted a recently developed 5’ RACE protocol^63^. Briefly, we extracted RNA from the head kidneys of WT and *rag2* mutant fish and performed cDNA amplification using a specific primer for the constant region. Template-switching further allowed us to amplify all *igh* recombination variants (**Figure 3c**, and see **Methods**).

This protocol produced a ∼600 base pair amplicon in WT fish, consistent with mature immunoglobulins^63^. Conversely, we did not detect these amplicons in *rag2* mutants (**Figure 3c**, bottom), and shorter faint bands are likely a result of non-specific amplification. This suggests the absence (or significant reduction) of V(D)J recombination in *rag2* mutants. These findings propose that the killifish *rag2* mutant is an immunodeficient genetic model.

### Functional characterization of *vgll3*-mediated melanoma in killifish

To explore the tumorigenicity of the melanocyte expansion, we first developed intramuscular transplantations. We initially used RFP^+^-derived^64^ primary killifish fibroblasts to facilitate visual confirmation of successful injections (**Figures S4a**, top). As expected from non-malignant cells, while the injected cells were detected in *rag2* mutants and WT controls at first, they did not persist (n=0/10 and n=0/8, respectively, see example in **Figures S4a**, bottom).

In contrast, cells isolated from a melanocyte expansion in the tail of *vgll3^Ex^*^1^*^/Ex^*^1^ mutants were successfully engrafted and generated an extensive melanoma-like tumor in *rag2* mutants (**Figure 4a**). As a control, these cells failed to engraft when transplanted into WT fish (0/3). Similarly, phenotypically normal melanocytes isolated from the tail of a young fish did not engraft into *rag2* mutants (0/3).

**Figure 4:**
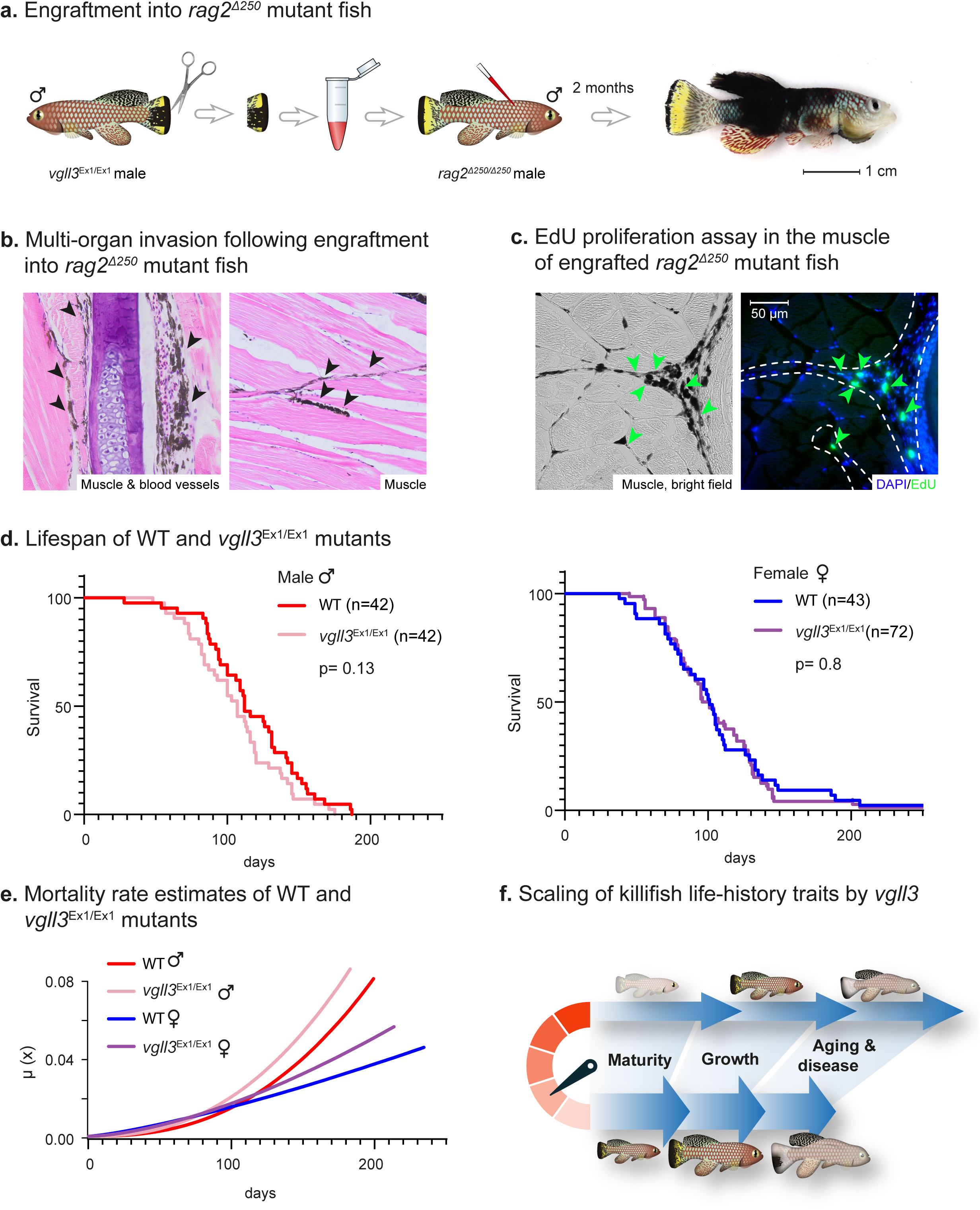
Age-related effects of *vgll3^Ex^*^1*/Ex*1^ mutants. **a)** Engraftment of primary cells derived from a melanocyte expansion in the tail of old *vgll3^Ex^*^1^*^/Ex^*^1^ mutant into a *rag2* mutant recipient results in a melanoma-like tumor. n = 3 biological replicates. **b)** H&E staining for the fish seen in (**a**) highlighting multi-organ invasion. Melanocytes are indicated with black arrows. **c)** Histological section from a fish engrafted with melanoma cells and injected with EdU. The slide was treated with H_2_O_2_ to remove the melanin pigment (right). Bright-field image of an adjacent section not treated with H_2_O_2,_ in which melanocytes are visible (left). The dashed line marks the corresponding area. Representative of n = 3 individuals. **d)** Lifespan of WT and *vgll3^Ex^*^1^*^/Ex^*^1^ fish, assessed separately for males (left) and females (right). P-values for differential survival in log-rank tests and fish numbers are indicated. **e)** Mortality curves as fitted by the simple Weibull model from survival data (see **Figure S4c**). **f)** A schematic model suggests that levels of the long isoform of VGLL3 can scale killifish life history, including puberty onset, growth rates, and age-related diseases.

Following successful engraftment, melanocytes were detected in multiple organs, invading skeletal muscle and blood vessels (**Figure 4b**). To visualize cell proliferation, tumor cells were engrafted into a new *rag2* recipient, and six weeks following engraftment the fish were injected with EdU (5-Ethynyl-2’-deoxyuridine, see **Methods**). Many EdU-positive nuclei were detected within the melanocyte-positive area (**Figure 4c**, and see quantification in **Figure S4b**).

Finally, while there was no overall difference in survival between WT and *vgll3^Ex^*^1*/Ex*1^ animals (**Figure 4d**, males: p=0.13, females: p=0.8, log-rank test), *vgll3^Ex^*1*^/Ex^*1 males had a faster increase in mortality hazards with age, also known as actuarial senescence^65^ (**Figure 4e**). We estimated age-specific mortality rates by a Bayesian survival trajectory analysis^66^, which identified the Weibull model as the best fit (**Figure S4c**, and see **Methods**). In the Weibull formula (µ_0_(x|b) = b0b1^b0^x^b0–1^), beta parameters describe either baseline mortality (b0) or mortality rate change with age (b1).

A substantial difference between groups can be then estimated using the Kullback–Leibler Divergence Calibration value^67,68^ (KLDC > 0.85), indicating that females display lower baseline mortality rates compared to males (**Figure S4d, e**, b0 KLDC: WT female vs. male=0.998; *vgll3^Ex^*^1*/Ex*1^ female vs. male=0.98). Strikingly, only *vgll3^Ex^*^1*/Ex*1^ males displayed a higher increase in age-related mortality (b1 KLDC: male *vgll3^Ex^*^1*/Ex*1^ vs. WT - 0.932, **Figures 4e, S4e,** and **Table S1**).

Together, these results identify *vgll3* as an antagonistic pleiotropic gene, which may act as a rheostat by scaling somatic growth, maturation, and age-related disease (**Figure 4f**). Additionally, our findings suggest distinct functional roles for specific *vgll3* isoforms, which may be evolutionarily conserved across vertebrates.

## Discussion

### Molecular mechanism of antagonistic pleiotropy in age-specific life histories

The antagonistic pleiotropy theory of aging maintains that aging evolves via the selection of genes that increase early-life performance at the cost of late-life performance. While experimental evolution and GWAS studies provide broad support for this theory, we often lack functional studies that explicitly link variation in individual genes to complex age-specific life histories.

In this study, we mutated two evolutionarily conserved isoforms of *vgll3* in the rapidly maturing turquoise killifish. Our data suggest that mutating the long isoform of *vgll3* by editing exon 1 results in a pleiotropic effect. On one hand, the mutants exhibited accelerated growth and maturity; on the other, they showed an increased incidence of age-related cancer and acceleration of mortality rate with age. Reducing the expression of both the short and long isoforms of *vgll3,* through a heterozygous mutation in exon 3, produced a similar effect on maturation.

However, disruption of both isoforms of *vgll3* in *vgll3^Ex^*^3*/Ex*3^ delayed maturation, indicating a dose- or isoform-dependent effect of *vgll3*. We identify a single gene that mediates antagonistically pleiotropic variation in complex, age-specific life-history traits in a vertebrate. These findings demonstrate the opposite trend of the extended lifespan and cancer protection observed in Laron Syndrome patients, a condition characterized by dwarfism caused by a growth hormone receptor mutation^69^.

Our findings demonstrate that specific mutations in *vgll3* enhance beneficial early-life effects, while being directly linked to detrimental late-life effects. Given the evolutionary conservation of *vgll3*’s function, this pleiotropic gene might be a broad regulator of maturity and aging across vertebrates.

### The role of *vgll3* in cancer

The *rag2* immunodeficient model facilitated the identification of melanoma-like cancer, which sporadically occurs during the aging of *vgll3^Ex^*^1*/Ex*1^ mutants. Yet, how *vgll3* can modulate cancer in killifish remains to be elucidated. Recent studies have implicated VGLL3 in tumor development^70,71^, either by promoting tumor cell proliferation and motility^71–73^, or conversely, as a tumor suppressor through interactions between the Hippo pathway and estrogen receptor alpha (ERα)^70^.

*Vgll3*, a member of the VGLL family, contains a conserved TONDU motif essential for binding with TEA domain-containing transcription factors (TEADs)^74^. It also features a unique glutamate-rich motif at the N-terminus, believed to mediate liquid-liquid phase separation^75^, and a histidine-rich motif at the C-terminus, associated with localization to nuclear speckles^76^.

Within the nucleus, a proposed mechanism could be via the role of VGLL3 in modulating chemosensitivity through promoting DNA double-strand break repair^77^. However, the role of VGLL3 in tumor development within the context of a healthy individual, and how it regulates maturity and growth, remains underexplored. Future studies applying RNA-seq and/or whole-genome sequencing of melanocytes derived from *vgll3* mutants could identify gene expression changes and potential additional mutations that contribute to tumorigenicity.

### The role of *vgll3* in maturity

What could be the underlying cause of the antagonistic pleiotropic phenotypes of *vgll3*? While *vgll3* encodes for ∼40% of maturation age variation in natural salmon populations^29,30^, the precise mechanisms by which *vgll3* functionally influences downstream gene regulation remain unknown^41^. Similarly, the sexually dimorphic nature of the effects of VGLL3^78–80^, such as female-biased autoimmune diseases, could also provide important insight.

An elegant study examining the selective forces shaping life-history trait evolution, explored the genomic basis of adaptations to seasonal habitat in 45 African killifish species^81^. Within their data, we identified *vgll3* as being positively selected in the short-lived African turquoise killifish (within the top 5% of all selected genes, p=4.9X10^-6^). Thus, suggesting the exciting possibility that this gene may have played a role in their adaptation to a naturally compressed lifespan.

Downstream to *vgll3*, several studies have begun to elucidate possible molecular mechanisms, such as involving the Hippo and TGF-β signaling pathways, and Sertoli cell function^40,41^. It was recently predicted that pleiotropic architectures might evolve more easily in transcriptional cofactors, such as *vgll3*^41^, which regulate genes through protein-protein interactions rather than direct DNA binding. The rationale behind this prediction is that cofactors can interact with multiple transcription factors, and simple changes, such as in expression levels, can orchestrate a strong effect on regulatory networks without compromising DNA binding.

Complex age-specific life histories are generally predicted to be highly polygenic. However, the AP theory postulates the existence of genes with broad pleiotropic effects across the life cycle. The developmental theory of aging, a physiological extension of AP, proposes that selection acting on early-life traits—from zygote to the onset of reproduction—can drive the evolution of aging. This occurs because genes beneficial during development may later become detrimental in adulthood.

Williams (1957)^3^ illustrated this concept with a hypothetical example of a gene involved in calcium metabolism in vertebrates: an allele that enhances bone calcification during development may later contribute to artery calcification, causing harm in old age. While it remains unclear whether calcium metabolism operates through such a mechanism, we identify *vgll3* as an antagonistically pleiotropic gene. This gene, involved in cell growth, promotes beneficial developmental traits but imposes late-life costs by increasing tumor incidence and actuarial senescence, ultimately shaping vertebrate life history.

## Figure legends

**Figure S1:**
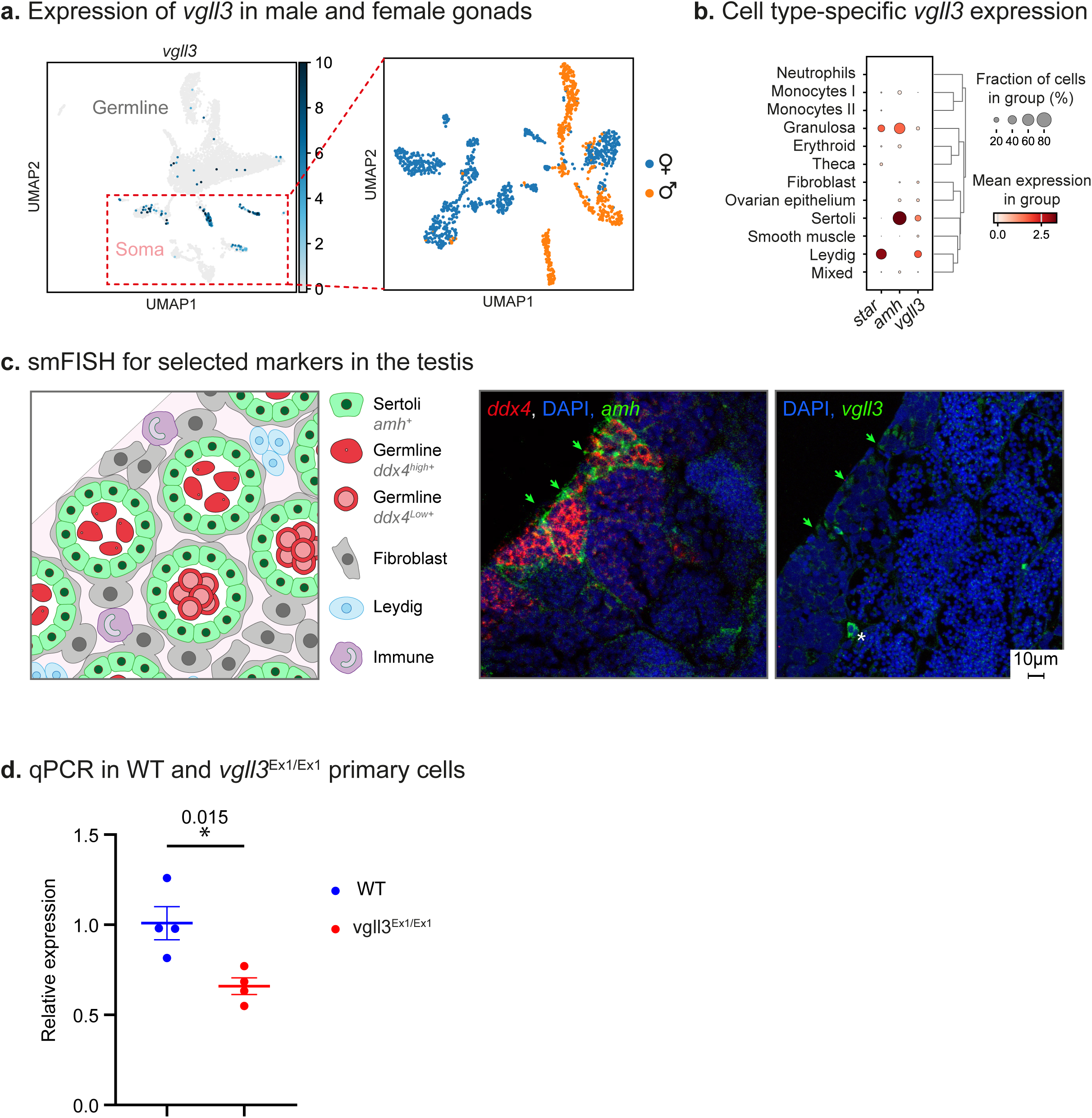
*vgll3* expression. **a)** UMAP analysis of 5,695 single cells from the testis and ovary of 1-month-old fish (data from^39^), suggesting that *vgll3* is primarily expressed in the somatic compartment (left). Somatic cells are clustered and color-coded by sex of origin (right). **b)** Dot plot representing the relative expression of selected cell-type-specific marker genes (according to^39^) and *vgll3*. **c)** Left: a model illustrating the spatial distribution of primary gonadal cell types in the killifish testis, including selected marker genes. For germ cells, *ddx4^high^*marks Spermatogonia, and *ddx4^low^* marks Spermatocytes. Adapted from^39^. Center: smFISH for germ cells (*ddx4,* red) and Sertoli cells (*amh*, green). Right: smFISH for *vgll3* (green). Representative of *n* ≥ 6 individuals. White asterisk marks autofluorescence from nucleated red blood cells. Scale bar: 10 µm. **d)** Relative *vgll3* expression measured using qPCR in primary fibroblast cultures of WT and *vgll3^Ex^*^1^*^/Ex^*^1^ males (see **Methods**). Error bars show mean ± SEM. Significance was measured using a two-sided Student’s t-test.

**Figure S2:**
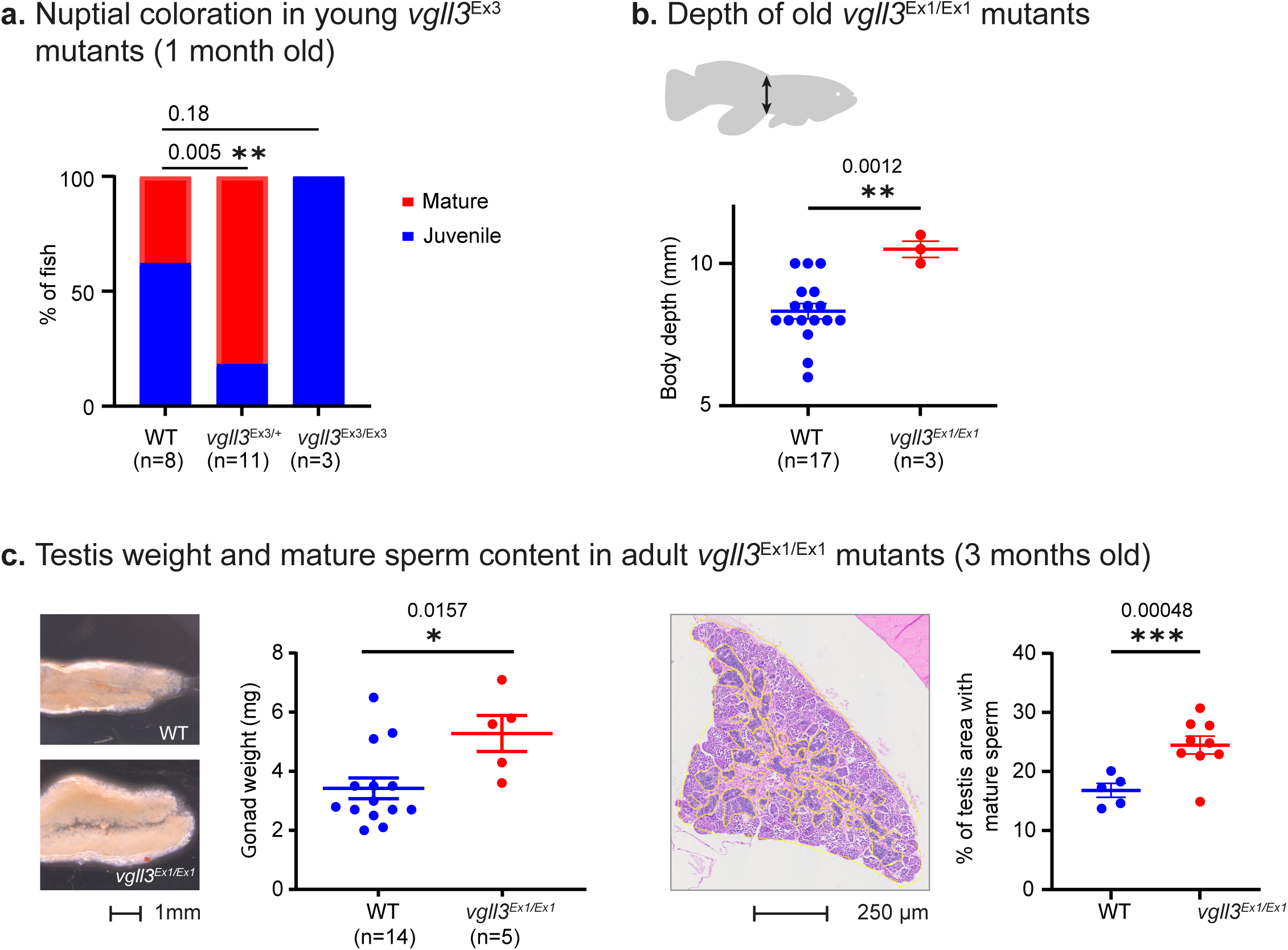
Gonadal effects of *vgll3* mutants. **a)** Proportion of fish displaying mature and juvenile coloration in 1-month-old WT, *vgll3^Ex^*^3^*^/+^*and *vgll3^Ex^*^3^*^/Ex^*^3^ male fish. Significance was measured using a χ^2^-test with the WT proportion as the expected value, and an FDR correction. **b)** Quantification of fish depth (distance between dorsal and anal fins). Error bars show mean ± SEM. Significance was measured using a two-sided Student’s t-test. **c)** Left: representative images of gonads dissected from 3-month-old WT and *vgll3^Ex^*^1^*^/Ex^*^1^ male fish. Center left: Quantification of gonad weight. Center right: representative image of an H&E staining of a testis, with Spermatozoa-containing cysts marked in yellow. scalebar: 250µm. Right: quantification of the relative area (%) containing spermatozoa in testis from three-month-old WT or *vgll3^Ex^*^1^*^/Ex1^*fish. At least 5 histological sections from ≥ 3 fish. Error bars show mean ± SEM. Significance was measured using a two-sided Student’s t-test.

**Figure S3:**
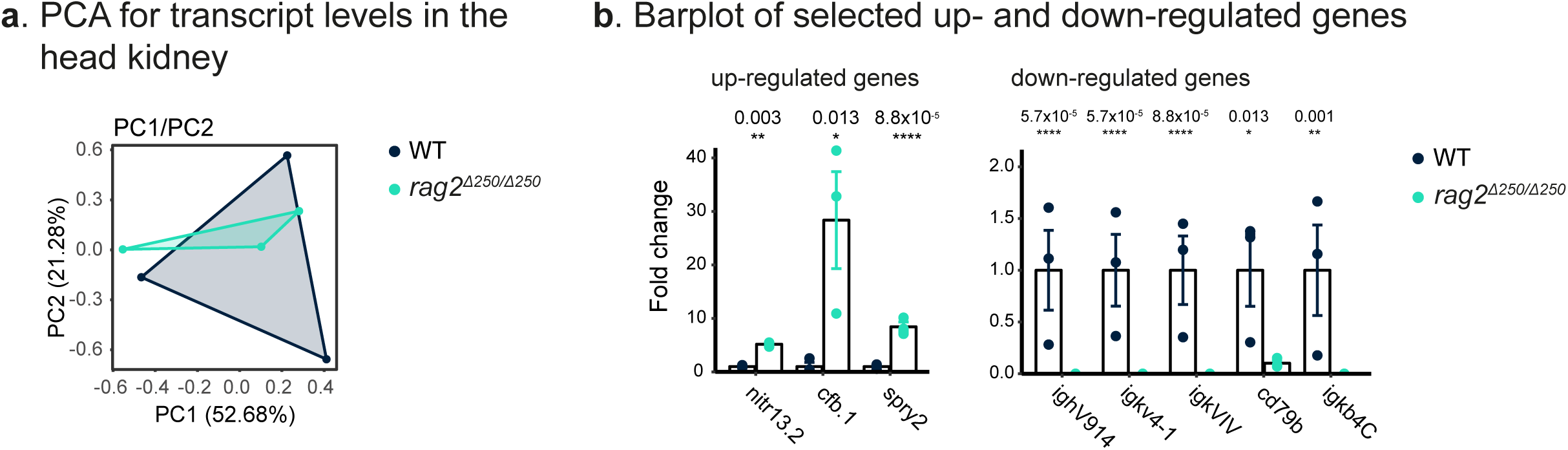
Altered immunity in *rag2* mutant fish. **a)** PC1/PC2 for transcript levels from the head kidney of WT (black) and *rag2* mutant fish (green). Each symbol represents an individual fish. n=3 for each genotype. **b)** Bar-plot of selected upregulated (left) and downregulated (right) genes, normalized to the WT. Error bars represent mean ± SEM. Significance was called as part of the edgeR pipeline. FDR-corrected p-values are indicated.

**Figure S4:**
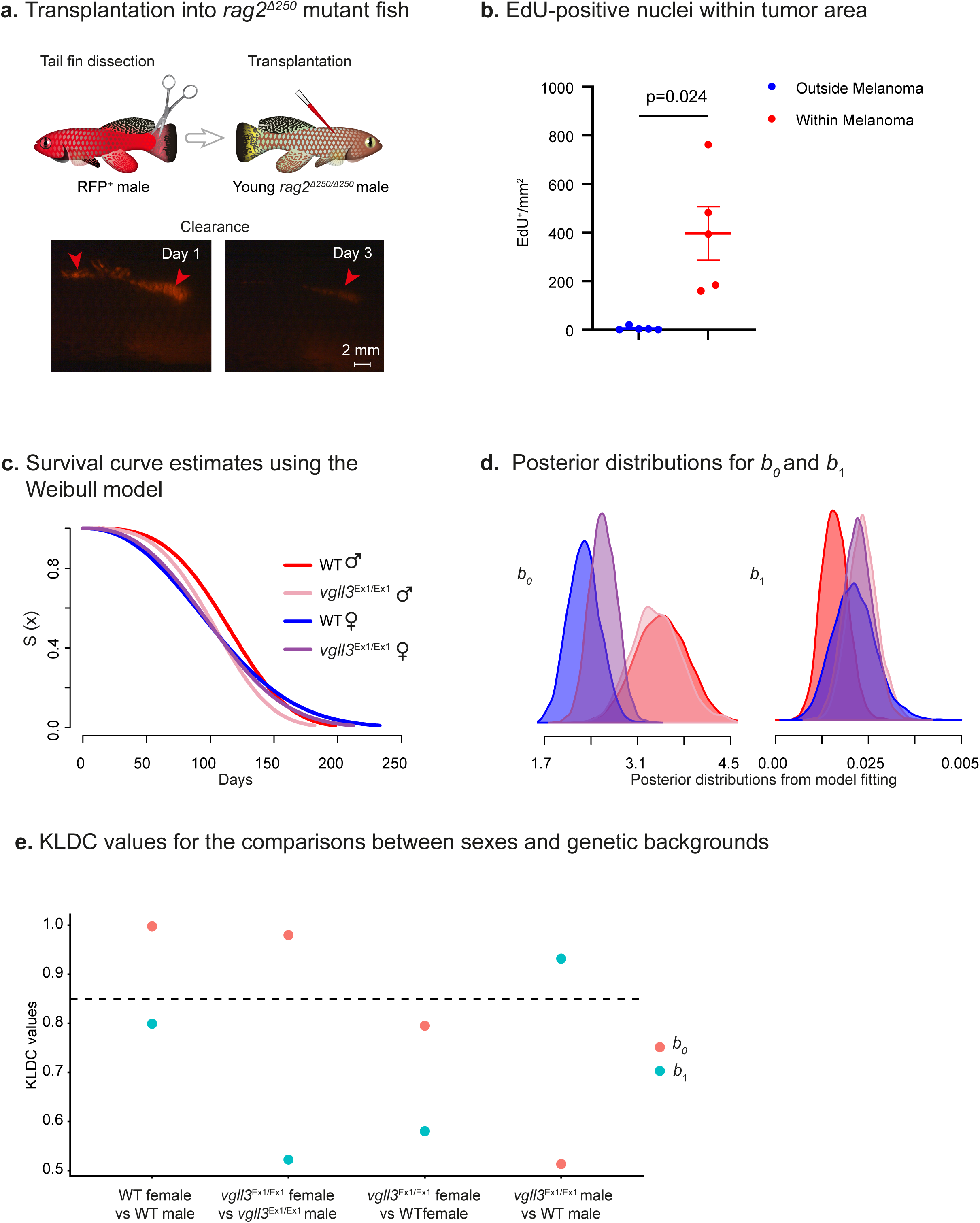
Transplantation assay and actuarial senescence. **a)** Injection of RFP^+^-derived primary cells into WT fish. n = 10 biological replicates. **b)** Quantification of density of EdU^+^ nuclei within and outside of the area populated by tumor-like cells, n= 5 images. Error bars represent mean ± SEM. Significance was measured using a two-sided paired Student’s t-test. **d)** The posterior distributions of b0 and b1 mortality parameters, which according to the Weibull formula (µ_0_(x|b) = b0b1^b0^x^b0–1^), describe either baseline mortality (b0) or mortality rate change with age (b1). **d-e)** The posterior distributions of mortality parameters (**d**) were compared between conditions using Kullback–Leibler divergence calibration (KLDC; **e**). KLDC values closer to 0.5 indicated minimal difference between the distributions and values approaching 1 implied a larger difference^82,83^. We considered a KLDC value > 0.85 to indicate a substantial difference between treatments^67,68^

## Acknowledgments

We thank the Harel lab for stimulating discussion and feedback on the manuscript. We thank Fatma Idrees and Reem Barakat for killifish maintenance. Supported by the ERC StG #101078188 (I.H.), Zuckerman Program (I.H.), ISF 2178/19 (I.H.), Israeli Ministry of Science 3-17631 (I.H.), 3-16872 (I.H.), the Moore Foundation GBMF9341 (I.H.), BSF-NSF 2020611 (I.H.), the Israeli Ministry of Agriculture 12-16-0010 (I.H), the Levi Eshkol scholarship of the Israeli Ministry of Science (E.M), the Czech Science Foundation (#22-01781O), and the Ministry of Education, Youth and Sports of the Czech Republic (#CZ.02.1.01/0.0/0.0/16_025/0007370) (R.F.), the Pamela and Paul Austin Research Center on Aging fellowship (T.A.).

## Author contributions

E.M., R.F., and I.H. designed the study. E.M., performed experiments with help from R.F., H.C. generated the *rag2* mutant. S.K. and H.C. assisted with maturity and lifespan experiments. T.A. designed and performed the analysis of the kidney RNA-seq of *rag2* mutants. All experiments were conducted under the supervision of I.H.. E.D. performed BaSTA analysis under the supervision of A.A.M.. E.M., R.F., T.A., and I.H. wrote the manuscript. All authors commented on the manuscript.

## Ethics declaration

The authors declare no competing interests.

## RESOURCE AVAILABILITY

### Material availability

All fish lines are available upon request.

### Data and Code availability

All raw RNA sequencing data, as well as processed datasets could be found in the GEO database, accession number GSE226279. The code is available in the GitHub repository for this paper https://github.com/Harel-lab/Killibow-rag2-mutant.

### Statistics and reproducibility

The number of biological replicates and statistical tests for each experiment are presented in the corresponding figure legends. All experiments were repeated independently with similar findings at least twice. Sample sizes are similar to those reported in previous publications^39,44^. This excludes the RNA-seq datasets and fish lifespan that had independent biological replicates. All fish were randomly assigned to each experimental group while controlling for age and genotype. Visual scoring of melanoma presence and size, maturity assessment and sperm content measurements were blindly and independently performed by two researchers. Power analysis was performed for melanoma experiments, predicting an increase of 50%of 50% in cancer incidence with an alpha of 0.05 and power of 80%:

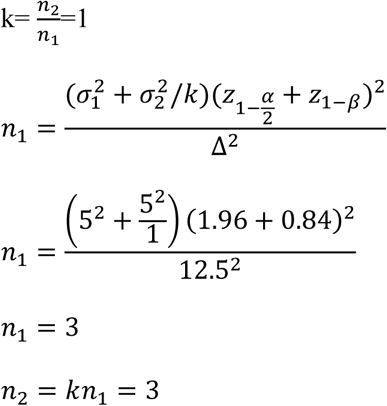

Fish were excluded from the lifespan cohort if they did not develop normally (e.g. did not inflate the swim bladder, “belly sliders”) or did not die a natural death (e.g. fell from the tank during routine maintenance). Gonads whose area was smaller than 25% of the maximal gonad area in their genotype were also excluded since they might be close to the end of the gonad and have non-representative cell-type compositions (see **Table S1**). These criteria were pre-established. Otherwise, no data was excluded. Data distribution was assumed to be normal in bench experiments because parameters measured, such as weight, percent proliferation, age at maturity and cancer incidence are usually normal. However, this was not formally tested. In RNA-seq analysis, the data were normalized, and the method of normalization is listed in the Methods section.

## EXPERIMENTAL MODEL AND SUBJECT DETAILS

### African turquoise killifish strain, husbandry, and maintenance

The African turquoise killifish (GRZ strain) were housed as previously described^39,42,44^. Fish were grown at 28 °C in a central filtration recirculating system with a 12 h light/dark cycle at the Hebrew University of Jerusalem (Aquazone ltd, Israel). Fish were fed once a day with live Artemia until the age of 2 weeks (#109448, Primo), and starting week 3, fish were fed three times a day on weekdays (and once a day on weekends), with GEMMA Micro 500 Fish Diet (Skretting Zebrafish, USA), supplemented with Artemia twice a day. All turquoise killifish care and uses were approved by the Subcommittee on Research Animal Care at the Hebrew University of Jerusalem (IACUC protocols #NS-18-15397-2, #NS-22-16915-3 and #HU-24-17607-4).

### CRISPR/Cas9 target prediction and gRNA synthesis

CRISPR/Cas9 genome-editing protocols were performed according to published protocols^42,44^. In brief, for gene targeting, a gRNA target site was identified using CHOPCHOP (https://chopchop.rc.fas.harvard.edu/)^84^, chosen gRNAs are listed below. Design of variable oligonucleotides, and hybridization with a universal reverse oligonucleotide was performed according to^42^, and the resulting products were used as a template for *in vitro* transcription. Each gRNA was *in-vitro* transcribed and purified using TranscriptAid T7 High Yield Transcription Kit (Thermo Scientific #K0441), according to the manufacturer’s protocol.

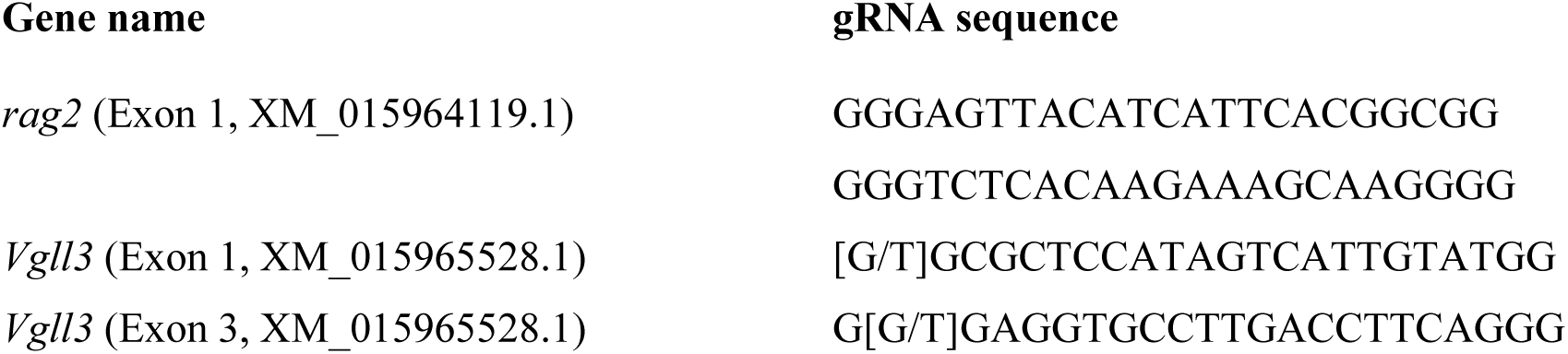

### Production of Cas9 mRNA

Experiments were performed according to^42,44,85^. The pCS2-nCas9n expression vector was used to produce *Cas9* mRNA (Addgene, #47929)^86^.Capped and polyadenylated mRNA was *in-vitro* transcribed and purified using the mMESSAGE mMACHINE SP6 ULTRA (ThermoFisher # AM1340).

### Microinjection of turquoise killifish embryos

For generation of the mutant fish using CRISPR/Cas9, microinjection of turquoise killifish embryos was performed according to^42^. Briefly, nCas9n-encoding mRNA (300 ng/μL) and gRNA (30 ng/μL) were mixed with phenol-red (P0290, Sigma-Aldrich) and co-injected into two-cell stage fish embryos. Sanger DNA sequencing was used for detecting successful germline transmission on F1 embryos using the following primer sequences: *rag2F*: TGATGTTTTCTGTTTTGACCCA and *rag2R*: TATCTGTGGGCAGGACCTGTA *Vgll3 exon1F:* GGACACAGCCCGTCTGAAGTCT *Vgll3 exon1R:* GAAGTTCGCCTCACCTGTTG *Vgll3 exon3F:* CTGTTCACCTACTTTAAGGGCG *Vgll3 exon3R:* TGTAGGTCCAGGTATCAGGGAG. We used several compound heterozygous mutations for Exon 3 indels, primarily *vgll3^ins^*^1^ and *vgll3^ins^*^5^ surrounding the PAM site.

### Maturity, growth and survival assays

### Maturity assessment

Since fish needed independent evaluation before genotyping, each fish was individually housed in a 1-liter tank starting at week 2. For assessing sexual maturity in males, the onset of tail coloration was visually scored at 30dph. Examples of juvenile and mature coloration can be found in **Figure 2a**.

### Growth and GSI measurements

Fish were euthanized with 500 mg/l tricaine (MS222, #A5040, Sigma), dried using Kimwipes and weighed. The testes were then dissected, washed in PBS, briefly dried using Kimwipes and weighed. GSI was calculated as follows: GSI = (gonad weight / total body weight) × 100. For depth measurement fish were imaged with a Canon EOS 250D digital camera. A ruler was included in each image to provide an accurate scale. Body depth was measured from the anterior end of the dorsal fin to the anterior end of anal fin. The depth was then calculated using ImageJ (1.52a), by converting pixel number to millimeters using the reference ruler.

### Melanoma incidence assessment

Male fish from both *vgll3^Ex^*^1*/Ex*1^ and WT backgrounds were routinely monitored starting at seven months of age. Fish in which the black stripe in the tail exceeded its normal boundaries and invaded into the yellow stripe in the tail were considered to exhibit melanoma-like expansion. Examples can be found in **Figure 2d**.

### Lifespan measurements

Constant housing parameters are very important for reproducible lifespan experiments^42,87^. After hatching, fish were raised with the following density control: up to 30 fish in a 1-liter tank for week 1, 5 fish in a 1-liter tank for week 2. Since these fish were also used for independent maturity assessment each fish was individually housed in a 1-liter tank starting at week 2. At the age of 4 weeks, adult fish were genotyped and housed individually in a 1-liter tank for the rest of their life. Both male and female fish were used for lifespan experiments and were treated identically. Fish mortality was documented daily starting at week 4. Lifespan analyses were performed using GraphPad Prism for all survival curves with a Kaplan-Meier estimator. A log-rank test was used to examine any significant differences between the overall survival curves of different experimental groups.

### Cell transplantation

For transplantation of fibroblasts, RFP^+^ cells^64^ were isolated from the tail, made into a cell line as described previously^44^ and were injected into *rag2* homozygous fish (40,000 cells/µl, at 4 µl per fish). For the injection, adult male recipient fish (∼1.5-month-old) were anesthetized, and intramuscular injection was performed using a Nanofil syringe (WPI, #NANOFIL).

For transplantation of melanoma-like cells, an 8-month-old *vgll3^Ex^*^1^*^/Ex^*^1^ mutant male with a melanocyte expansion in the tail fin was euthanized, part of the fin was dissected, finely minced with scissors and digested with 0.2% Collagenase Type P (Merck Millipore) and 0.12% Dispase II (Sigma Aldrich) in Leibovitz’s L-15 Medium (Gibco) for 30 min at RT. Cells were filtered using a Falcon 40 µm filter, centrifuged at 0.4g for 10 min and resuspended in L15 with 10% FBS and injected dorsally in *rag2* and WT recipients. Following *in-vivo* expansion of the melanoma cells in the *rag2* recipients, dorsal muscle tissue containing melanoma was dissected and prepared for transplantation into recipients again as described above. Fish were monitored following engraftment for ethical purposes, and melanoma development was visually confirmed.

### Hematoxylin and eosin

Tissues samples were processed as described previously^39,42,94–100,44,85,88–93^. Briefly, fish were euthanized with 500 mg/l tricaine (MS222, #A5040, Sigma). The body cavity of the fish was opened and fixed for 72 h in 4% PFA solution at 4 °C. Samples were dehydrated and embedded in paraffin using standard procedures. Sections of 5-10 μm were stained with Hematoxylin and Eosin and examined by microscopy. A fully motorized Olympus IX83 microscope with an Olympus DP28 camera was used to collect images. The total area and the area containing Spermatozoa in images of three-month-old WT and *vgll3^Ex^*^1*/Ex*1^ mutant testes was calculated using ImageJ. Testes that had a total area that was smaller than 25% of the maximal area measured for that genotype were censored since they might be close to the end of the testes and have non-representative cell-type compositions (see **Table S1**).

### EdU staining

A male *rag2* mutant fish that has been injected with melanoma, received an IP injection of EdU (50mg/kg in DMSO). After 6 hours the fish was sacrificed, its trunk was dissected, fixed for 48 h in 4% formaldehyde at 4 °C, dehydrated in an ethanol and chloroform series, embedded in paraffin, and sectioned. Consecutive 7 μm sections were collected individually on slides, deparaffinized and rehydrated. One section was immediately mounted with Fluoroshield containing DAPI (Sigma-Aldrich # F6182). An adjacent section was treated overnight with 30% H_2_O_2_ to bleach the pigment of the melanoma (according to^101^), and then stained with the EdU-Click Kit and mounted with Fluoroshield containing DAPI. Imaging was performed as described above and analyzed in ImageJ.

### Single molecule FISH

smFISH was performed as described previously^39^. In brief, split initiator hybridization probes for FISH chain reaction (HCR version 3.0) were designed and manufactured by Molecular Instruments for the following genes: *amh* (<u>XM_015977279.1</u>, *ddx4* (<u>XM_015957842.1</u>) and *vgll3* <u>(XM_015965528.2).</u> Then, 10-µm paraffin slices (as described above) were baked for 1 h at 60 °C, and smFISH was performed according to the manufacturer’s guidelines. In brief, after rehydration, slides were boiled for 15 min in 0.01 M citrate buffer (Sigma-Aldritch, C8532) and permeabilized with 20 µg ml^−1^ proteinase K (A&A Biotechnology, 1019-20-5) in PBS for 15 min at 37 °C, before being washed with PBS. Slides were pre-hybridized in hybridization buffer provided by the manufacturer (Molecular Instruments) and then hybridized with the indicated probe (20 nM, in hybridization buffer) at 37 °C overnight. After washing, the signal was amplified with either custom-made green (488) or red (546) fluorophores at room temperature overnight. Slides were washed with 5×SCCT, and autofluorescence was quenched using a TrueVIEW Autofluorescence Quenching Kit (Vector Labs, SP8500) according to the manufacturer’s protocol. Slides were mounted with VECTASHIELD containing DAPI (Vector Labs, 30326), and images were collected with an Olympus FV-1200 confocal microscope and processed in ImageJ software.

### Profiling of V(D)J recombination

Amplification of the V(D)J locus was performed according to^63^, with slight modifications. RNA was collected from the kidneys of 3 WT and 3 *rag2* mutant males and purified using the Direct-zol RNA Miniprep kit (Zymo Research, #R5052). Forty-five ng of RNA was combined with 2 μl of 10 μM gene-specific primer (GSP, **Table S1**), homologous with the second constant-region exon of N. furzeri IgHM. The reaction volume was brought to a total of 8 μl with nuclease-free water, and the resulting mixture was incubated for 2 min at 70 °C to denature the RNA, then cooled to 42 °C to anneal the GSP^102^.

Following annealing, the RNA-primer mixture was combined with 12 μl of reverse transcription master-mix (SMARTScribe Reverse Transcriptase, Takara-Clontech #639538) including the reverse-transcriptase enzyme and template-switch adapter SmartNNNa primer (**Table S1**). This primer consists of a constant sequence used for further PCR steps, a variable sequence containing multiple random nucleotides (N) that function as a unique molecular identifier (UMI) and several deoxyuridine bases (U), which are used to degrade the primer following amplification and a 3’-terminal sequence of riboguanosine residues, which anneal to cytidine residues added by the reverse transcriptase and enable template switching (See **Figure 3c**). The complete reaction mixture was incubated for 1 h at 42 °C for the reverse transcription reaction, then mixed with 2 μl of uracil DNA glycosylase (NEB, #M0280S, 1000 units) and incubated for a further 40 min at 37 °C to digest the template-switch adapter.

Following reverse transcription, the reaction product was purified using a standard PCR purification kit (QIAGEN #28104). The reaction mixture then underwent PCR amplification with the PCRBio HS Taq Mix Red (#PB10.23) using the IGH-B and M1SS primers (**Table S1**), according to the manufacturer’s instructions and ran on a 1% agarose gel.

### RNA-seq library preparation

Head kidney was isolated as described previously^42^. Samples were disrupted by bead beating in 300 μl of TriZol (Sigma) and a single 3 mm metal bead (Eldan, BL6693003000) using TissueLyzer LT (QIAGEN, #85600) with a dedicated adaptor (QIAGEN, #69980). Beating was performed at 50 Hz for 5 min. RNA extraction was performed with TRI Reagent (Sigma, #T9424) RNA concentration and quality were determined by using an Agilent 2100 bioanalyzer (Agilent Technologies). Library preparation was performed using KAPA mRNA HyperPrep Kit (ROCHE-08105936001) according to the recommended protocols. Library quantity and pooling were measured by Qubit (dsDNA HS, Q32854), size selection at 4% agarose gel. Library quality was measured by Tape Station (HS, 5067-5584). Libraries were sequenced by NextSeq 2000 P2, 100 cycle,75 bp single-end (Illumina, 20024906) with ∼40 million reads per sample.

### RNA sequencing analysis

Quality control and adapter trimming of the fastq sequence files were performed with FastQC (v0.11.8)^103^, fastx-toolkits (v0.0.13), Trim Galore! (v0.6.4)^104^, and Cutadapt (3.4)^105^. Options were set to remove Illumina TruSeq adapters and end sequences to retain high-quality bases with *phred* score > 20 and a remaining length > 20 bp. Successful processing was verified by re-running FastQC. Reads were mapped and quantified to the killifish genome Nfu_20140520^98,106^ using STAR 2.7.6a^107^. Differential gene expression between *rag2* mutant and WT fish was performed using the edgeR package, (v3.32.1)^108,109^ by the classic model (exactTest function).

### Gene Ontology Enrichment Analysis

Enriched Gene Ontology (GO) terms associated with transcripts level was identified using GO implemented in R package clusterProfiler (v3.18.1)^110^ with the threshold of FDR < 0.05 and log2(FC) > 1. GO terms were based on human GO annotations from org.Hs.eg.db (v3.13.0)^111^ and AnnotationDbi (v1.54.1)^112^.

### Generation of a *vgll3^Ex^*^1/Ex^^1^ primary cell culture

Cells were isolated as described previously^44^. Briefly, adult fish were sedated with MS-222 (200 mg/L Tricaine), and a 2–3 mm piece of their tail fin was trimmed using a sterile razor blade. Following disinfection with 70% ethanol, tissue samples were incubated for 2 h with 1 mL of an antibiotic solution containing Gentamicin (50 µg/mL Gibco) and Primocin^TM^ (50 µg/mL, InvivoGen) in PBS at room temperature. Tissues were then transferred into an enzymatic digestion buffer (200 µL, in a 24-well plate) containing Dispase II (2 mg/mL, Sigma Aldrich) and Collagenase Type P (0.4 mg/mL, Merck Millipore) in Leibovitz’s L-15 Medium (Gibco), minced with a sterile pair of scissors and incubated for 15 min. The digested tissue was then mixed with 400 µL of complete Leibovitz’s L-15 growth medium (Gibco), supplemented with Fetal Bovine Serum (15% FBS, Gibco), penicillin/streptomycin (50 U/ml, Gibco), Gentamicin (Gibco, 50 µg/ml) and Primocin^TM^ (50 µg/ml, InvivoGen). On the following day, media was carefully replaced, and during the first 7 days, cells were daily washed with fresh media before adding new media. When cells reached 85–90% confluency, they were passaged with Trypsin-EDTA 0.05% (0.25% Trypsin-EDTA, diluted in PBS). Cells were incubated in a 28 °C humidified incubator (Binder, Thermo Scientific) with normal air composition.

### Quantitative PCR

In order to measure *vgll3* expression on mutants, we performed real time qPCR using RNA extracted from cultured fibroblasts from WT and *vgll3^Ex^*^1^*^/Ex^*^1^ mutant fish that were treated with Etoposide (Sigma, E1383) at 50uM for 48 h to induce *vgll3* expression^77^. Cells were then washed with PBS and cultured with fresh culture media for 10 more days. RNA was extracted using TRI reagent (Sigma-Aldrich, T9424) and purified using a Direct-zol RNA Miniprep Kit (Zymo Research, R2052) according to the manufacturer instructions. cDNA was prepared using a Verso cDNA Synthesis Kit (Thermo Scientific # AB1453A). qPCR was performed using Fast SYBR Green Master Mix (ThermoFisher scientific # 4385612) on a QuantStudio5 Real-Time PCR System (ABI #A34322) using the following primers:

*vgll3*(Ex3)*-F*: GCTAACTATTATATGGAAAGGTGGTGG,

*vgll3*(Ex3)*-R*: GAGTTGGCTTGAGAGGGAAAG,

*actin-F*: ATGTTTGAGACCTTCAACACACC,

*actin-R*: TCCATCACGATACCTGTGGTTC.

*tbp-F*: CGGTTGGAGGGTTTAGTCCT

*tbp-R*: GCAAGACGATTCTGGGTTTG

*insr-F*: TGCCTCTTCAAACCCTGAGT

*insr-R*: AGGATGGCGATCTTATCACG.

### BaSTA Analysis

We used Bayesian survival trajectory analysis in the BaSTA package^66^ (ver. 1.9.5) to estimate age-specific mortality rates. We compared ten models of age-specific mortality rates using Gompertz, Logistic, and Weibull mortality functions, with either a simple, Makeham^113^ or bathtub shape^114^, or an exponential model with a simple shape, representing no senescence. We ran four parallel simulations of each model, for 600,000 iterations, with a burn-in of 60,001 iterations and a thinning of 600, allowing for robust convergence of all models in the ‘multibasta’ function of the BaSTA package. Models were then compared using deviance information criteria (DIC values^115^).

The best-fitting model with the lowest DIC) was identified as the Weibull model with simple shape parameters (see **Table S1**). Colchero et al. 2012^66^ recommended the addition of a Makeham constant term (c), representing age-independent mortality, to avoid potentially immortal individuals. Therefore, we compared outputs between Weibull models with simple versus Makeham shape parameters. The model parameters did not converge for the Weibull Makeham model with this dataset, even when it was run for 1,000,000 iterations, so we proceeded with the simple Weibull model: µ_0_(x|b) = b0b1^b0^x^b0–1^.

Here, beta parameters describe baseline (b0) and increase in mortality with age following a power function (b1). To achieve low serial autocorrelation for all parameter estimates (< 5%), we ran the simple Weibull model for 1,000,000 iterations, with a burn-in of 100,001 iterations and thinning of 1000 (see **Table S1**). The posterior distributions of all mortality parameters converged and were compared between treatments using Kullback–Leibler divergence calibration (KLDC; **Figure S4e**). We considered a KLDC value > 0.85 to indicate a substantial difference between treatments, following established practice^67,68^.

